# Diverse chemical compounds target *Plasmodium falciparum* plasma membrane lipid homeostasis

**DOI:** 10.1101/444158

**Authors:** Suyash Bhatnagar, Sezin Nicklas, Joanne M. Morrisey, Daniel E. Goldberg, Akhil B. Vaidya

**Affiliations:** Center for Molecular Parasitology, Department of Microbiology and Immunology, Drexel University College of Medicine, Philadelphia, Pennsylvania, United States of America; Department of Medicine, Division of Infectious Diseases, Washington University School of Medicine, St. Louis, Missouri, United States of America

## Abstract

Lipid homeostasis is essential for the maintenance of life. We previously reported that disruptions of the parasite Na^+^ homeostasis via inhibition of PfATP4 resulted in elevated cholesterol within the parasite plasma membrane as assessed by saponin sensitivity. A large number of compounds have been shown to target the parasite Na^+^ homeostasis. We therefore screened the same collection of 800 compounds to identify chemotypes that disrupted the parasite plasma membrane lipid homeostasis. Here, we show that the compounds disrupting parasite Na^+^ homeostasis also induced saponin sensitivity, an indication of parasite lipid homeostasis disruption. Remarkably, 13 compounds were identified that altered plasma membrane lipid composition independent of Na^+^ homeostasis disruption. Further studies suggest that these compounds target the *Plasmodium falciparum* Niemann-Pick Type C1-Related (PfNCR1) protein, which is hypothesized to be involved in maintaining plasma membrane lipid composition. PfNCR1, like PfATP4, appears to be targeted by multiple chemotypes with potential for drug discovery.

## Introduction

Malaria is a major heath burden on developing countries with billions of people at risk of contracting the disease. The World Health Organization (WHO) estimated that 148-304 million cases of malaria cases occurred in 2015, with 235,000-639,000 deaths resulting from the disease ^1^. The rise in resistance to frontline therapies highlights the need for the identification of novel targets and therapeutics ^2-3^. Recent investigations have led to the identification of three chemically distinct series of compounds (pyrazoleamides ^4^, spiroindolones ^5^ and dihydroisoquinolones (DHIQ) ^6^) that inhibit the parasite’s Na^+^/H^+^ antiporter PfATP4 resulting in the rapid influx of Na^+^ and simultaneous alkalinization of the cytoplasm ^7^.

Saponin has been long used for permeabilizing cholesterol-rich mammalian cells, including erythrocytes ^8^. During intraerythrocytic development of the parasite, the parasite plasma membrane (PPM) contains very low levels of cholesterol ^9-10^, rendering the parasites resistant to permeabilization by the cholesterol-dependent detergent, saponin. This differential cholesterol content between host and parasites plasma membranes has been exploited to isolate malaria parasites from their host red blood cells (RBCs). We previously reported that Na^+^ influx either via PfATP4 inhibition or a Na^+^ ionophore led to rapid alterations in the lipid composition of the parasite plasma membrane (PPM) and acquisition of cholesterol, as judged by saponin sensitivity ^11^. This effect was reversible, suggesting an active process involved in maintenance of low cholesterol levels in the PPM ^11^.

We also found that inhibition of a PPM protein we termed *Plasmodium falciparum* Niemann-Pick Type C1-Related Protein (PfNCR1) ^12-13^, caused a rapid alterations to PPM lipid composition ^13^. However, this alteration of cholesterol distribution through the inhibition of PfNCR1 was achieved without disrupting the Na^+^ homeostasis of the parasite ^13^. These results suggest two different targets in the PPM. Inhibition of either leads to rapid changes in lipid composition of the PPM as revealed by reversible acquisition of saponin-sensitivity, one dependent (PfATP4) on and the other independent (PfNCR1) of Na^+^ influx into the parasite.

The Medicine for Malaria Venture (MMV) has made available a collection of compounds to aid drug discovery. The first collection named the Malaria Box consisted of compounds representing a diverse chemical series that were identified from phenotypic screens against *P. falciparum* ^14^. The success of the Malaria Box was followed by the release of a second collection called the Pathogen Box. The Pathogen Box consists of drug like compounds that were active against a variety of pathogens (http://www.pathogenbox.org/) including *P. falciparum*. Both collections of compounds have been screened by the Kirk group for their ability to target the Na^+^ homeostasis of the parasite ^15-16^. These studies revealed that widely diverse chemical classes possessed the ability to disrupt the parasite Na^+^ homeostasis ^15-16^. A chemogenetic screen of the Malaria Box revealed multiple druggable targets in the *P. falciparum* genome ^12^.

We have devised an assay for rapid determination of lipid composition alterations of the PPM and have applied it to screen both the Malaria and Pathogen Boxes. This screen showed that compounds inhibiting PfATP4 also induced lipid composition changes within the PPM. In addition, the screen also identified compounds that appear to act by inhibition of PfNCR1, thus providing hits for future exploration of chemotypes targeting another PPM resident protein.

## Results

### Establishing a high-throughput assay

In our previous study, saponin mediated PPM permeabilization was assessed by monitoring the loss of cytosolic aldolase in western blots probed with anti-aldolase antibody ^11^. This method limited the rate at which large number of compounds could be processed. Thus, we employed parasites that stably express yeast dihydroorotate dehydrogenase (yDHODH) fused to green fluorescent protein (GFP) in the cytosol ^17-18^. A spectrofluorometer was used to record GFP emissions from these parasites to represent the cytoplasmic content of the parasites. Because different batches of saponin have varying concentrations of sapogenin, the component responsible for cholesterol-dependent permeabilization, we standardized the concentration of saponin used for the screen. Parasites were treated for 2 h with either vehicle (DMSO) or the pyrazoleamide PA21A092. Treated parasites were subsequently exposed to incremental concentrations of saponin to establish the concentration that displayed differential saponin sensitivity. Saponin exposure was conducted in Albumax-free RPMI to avoid cholesterol from Albumax confounding saponin sensitivity. Based on this the saponin concentration of 0.08% w/v was used for the remainder of the assays (Figure 1A). We further validated the assay by using other compounds known to induce saponin sensitivity. We assessed the dose dependency for the pyrazoleamide PA21A050, spiroindolone KAE609 and the Na^+^ ionophore Maduramicin. Similar to previous results ^11^, we observed a dose-dependent loss of GFP in drug-treated parasites due to saponin induced permeabilization (Figure 1B). Effective concentrations for 50% loss of cytosolic GFP (EC_50_) for PA21A050 and KAE609 were similar to the EC_50_ values observed for parasite growth inhibition ^4-5^.

**Figure 1:**
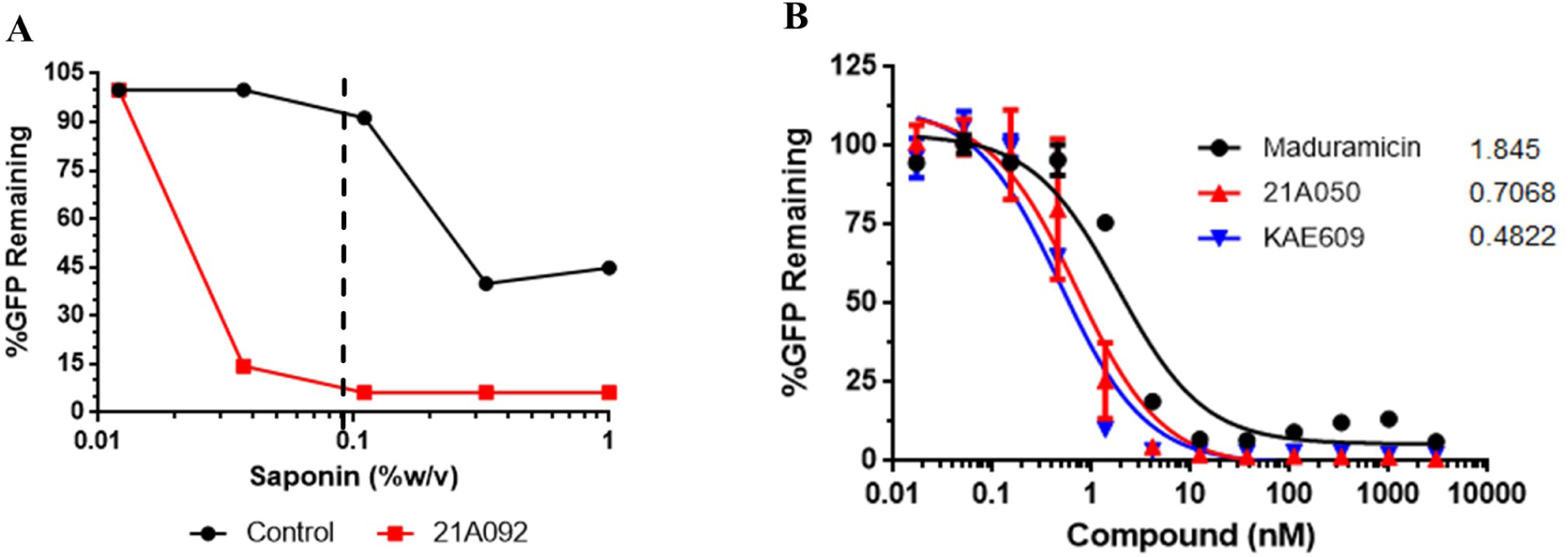
Assessment of differential saponin sensitivity. (A) Trophozoite stage NF45 yDHODH-GFP parasites treated for 2 h with vehicle control (Black) or 100nM pyrazoleamide 21A092 (Red). Parasite were released from host cells using a final saponin concentration of 1%, 0.33%, 0.11%, 0.037% or 0.012% w/v and GFP emission were recorded (n=2). (B) Trophozoite stage NF54 yDHODH-GFP were treated for 2 h with indicated concentrations of the Na^+^ ionophore Maduramicin (Black), pyrazoleamide PA21A050 (Red) and spiroindolone KAE609 (Blue). GFP emissions were plotted from parasites released using 0.08% saponin w/v. (n=2). Values adjacent to curves represent IC_50_ (nM).

### Screening for inhibitors of parasite cholesterol homeostasis

The screen was designed to identify rapid disruptors of PPM lipid homeostasis by assessing saponin sensitivity after a 2 h treatment. All compounds were diluted from stock concentration of 10mM to 100μM in DMSO. The compounds were used at a final concentration of 1μM thereby ensuring 1% DMSO in all wells. Synchronized trophozoite stage parasites were plated at 5% hematocrit in 96 well plates and treated for 2 h with 1μM of each compounds. Parasites were subsequently exposed to 0.08% w/v saponin in Albumax-free RPMI1640 and collected by centrifugation. The saponin treated parasite pellets were then lysed in radioimmunoprecipitation assay (RIPA) buffer to release parasite proteins. A spectrofluorometer was used to record GFP emissions at 510nM with an exciting wavelength of 488nm (Figure 2).

**Figure 2:**
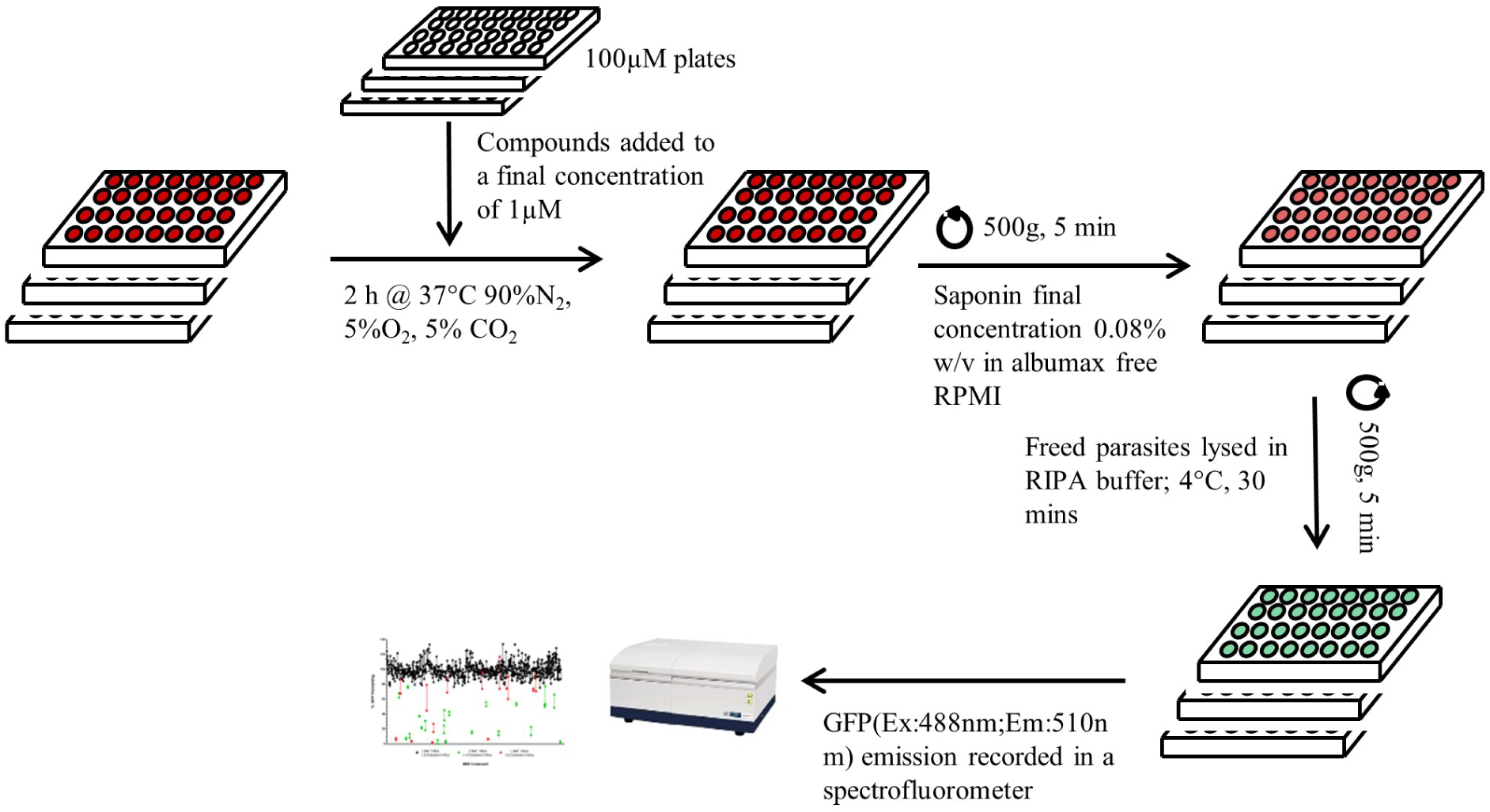
Schematic for screening compounds. Trophozoite NF54 yDHOD-GFP parasites were plated in 96 well plates with 1μM of the compounds and placed in culture conditions for 2 h. Parasites were isolated from host RBCs by exposing cultures in 0.08% w/v saponin prepared in Albumax-free RPMI1640 followed by centrifugation. Isolated parasites were washed 2x with Albumax supplemented RPMI1640. Parasite pellets were lysed in RIPA buffer at 4°C for 30 min and placed in a spectrofluorometer to record GFP emission at 510nm with a 488nm excitation.

The screen of the Malaria Box identified 22 compounds that showed a significant reduction in GFP content as a measure of saponin sensitivity compared to parasites treated with the combination of 100nM Atovoquone (Atv) and 1μM Proguanil (Pro) (Figure 3A; Table 1). The combination Atv/Pro was used as it has been previously reported to be lethal to parasites expressing yDHODH ^17^, but demonstrate no saponin sensitivity. Using the same parameters, the Pathogen Box compounds were also screened, leading to identification of an additional 19 compounds that induced saponin sensitivity after a 2 h exposure to 1μM of the compounds (Figure 3B; Table 2). The Malaria Box ^15^ and Pathogen Box ^16^ have been screened previously by the Kirk laboratory for compounds that disrupt Na^+^ homeostasis in *P. falciparum*. These screens identified 16 compounds from the Malaria Box and 12 compounds from the Pathogen Box as causing Na^+^ influx into parasites when used at 1 μM concentrations. We compared the compounds identified in our screen for saponin sensitivity induction with the Na^+^ influx induction data from the Kirk laboratory. Based on this comparison, we found compounds that induced both Na^+^ influx and saponin sensitivity (Table 1, Table 2). We found 16/28 compounds from the Malaria Box and 10/11 compounds from the Pathogen Box that were identified in the Na^+^ influx screen and displayed significantly saponin sensitivity. These results support the hypothesis that Na^+^ homeostasis disruption leads to altered lipid homeostasis of the PPM. Examination of the list of compounds identified by the Kirk laboratory as Na^+^ influx active revealed 15 compounds that induced saponin sensitivity independent of Na^+^ influx (Table 1, Table 2). These compounds appeared to target a previously unexplored pathway that maintained the plasma membrane lipid homeostasis.

**Figure 3:**
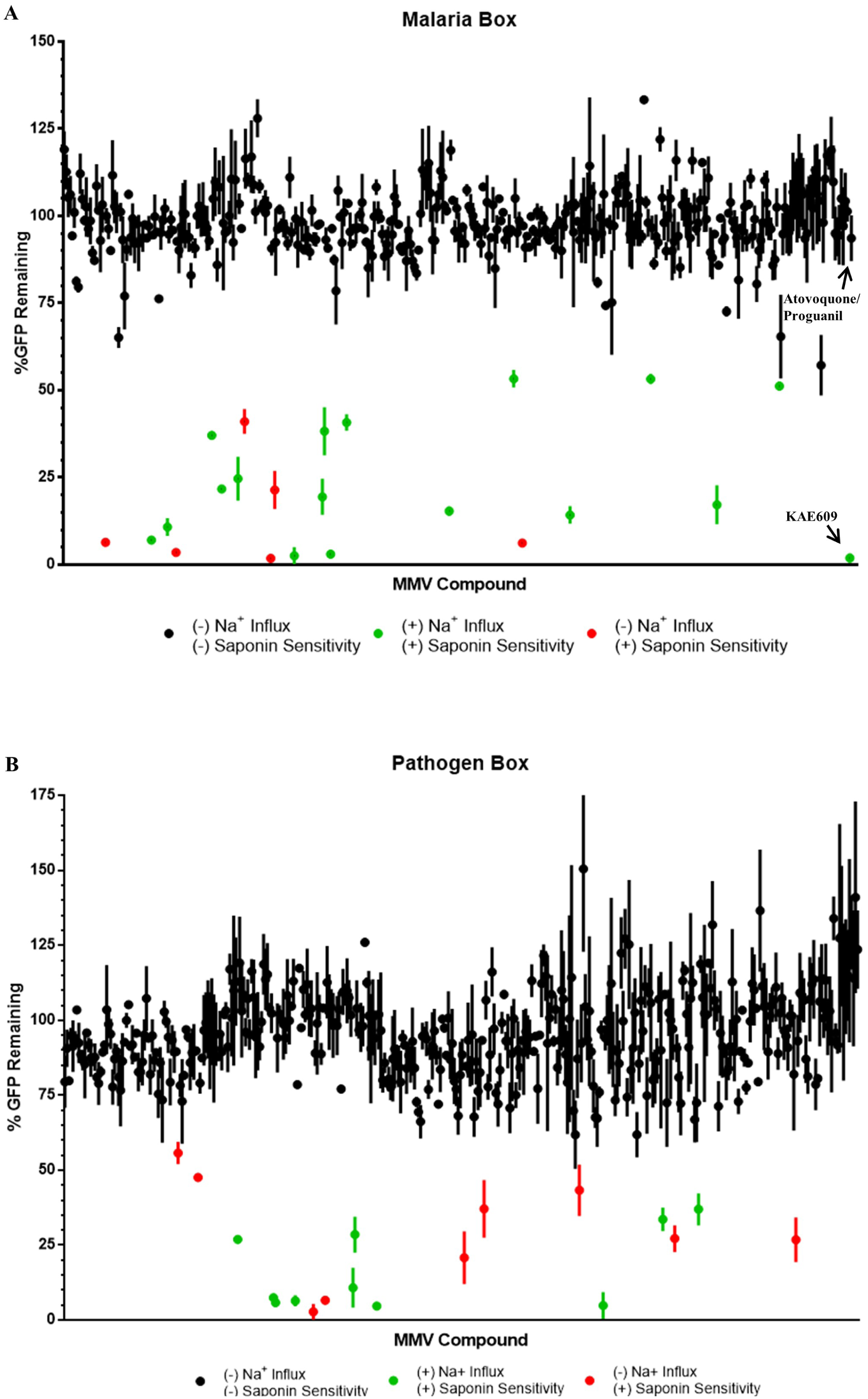
Compounds screened for induced saponin sensitivity. Parasite GFP content represented in % after 2 h treatment with each of the compound in (A) the Malaria Box and (B) the Pathogen Box. Black circles indicate no significant saponin sensitivity, green circles represent saponin sensitivity induced by Na^+^ influx and red circles display saponin sensitivity with no changes in cytoplasmic [Na^+^]. Na^+^ homeostasis properties of the compounds were based on published reports from the Kirk laboratory ^15-16^. The spiroindolone KAE609 was the positive control for saponin sensitivity and 100nM Atovoquone/1μM Proguanil served as the negative control. (p < 0.05)

**Table 1:**
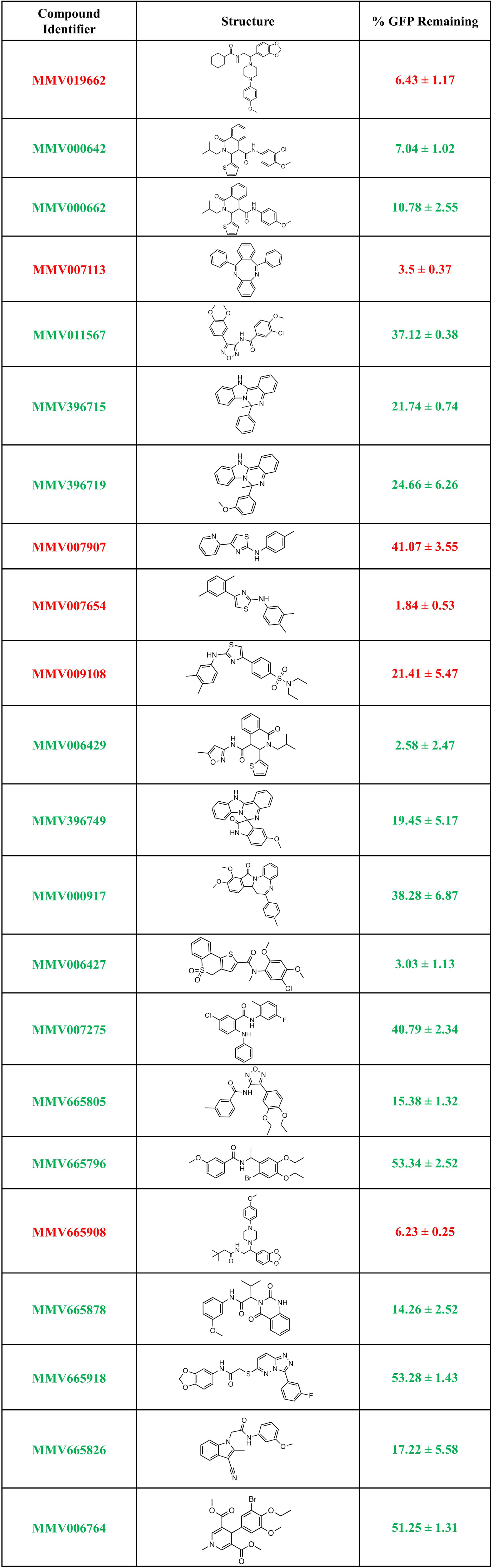
Candidates from the Malaria Box disrupting lipid homeostasis. Compounds from the Malaria Box that were designated as candidates inducing saponin sensitivity into the parasite plasma membrane (mean ± SEM from 2 independent experiments). Compounds in Green represent Na^+^-dependent and Red represent Na^+^-independent saponin sensitivity.

| Compound Identifier | Structure | % GFP Remaining |
| --- | --- | --- |
| MMV019662 |  | 6.43 ± 1.17 |
| MMV000642 |  | 7.04 ± 1.02 |
| MMV000662 |  | 10.78 ± 2.55 |
| MMV007113 |  | 3.5 ± 0.37 |
| MMV011567 |  | 37.12 ± 0.38 |
| MMV396715 |  | 21.74 ± 0.74 |
| MMV396719 |  | 24.66 ± 6.26 |
| MMV007907 |  | 41.07 ± 3.55 |
| MMV007654 |  | 1.84 ± 0.53 |
| MMV009108 |  | 21.41 ± 5.47 |
| MMV006429 |  | 2.58 ± 2.47 |
| MMV396749 |  | 19.45 ± 5.17 |
| MMV000917 |  | 38.28 ± 6.87 |
| MMV006427 |  | 3.03 ± 1.13 |
| MMV007275 |  | 40.79 ± 2.34 |
| MMV665805 |  | 15.38 ± 1.32 |
| MMV665796 |  | 53.34 ± 2.52 |
| MMV665908 |  | 6.23 ± 0.25 |
| MMV665878 |  | 14.26 ± 2.52 |
| MMV665918 |  | 53.28 ± 1.43 |
| MMV665826 |  | 17.22 ± 5.58 |
| MMV006764 |  | 51.25 ± 1.31 |

**Table 2:**
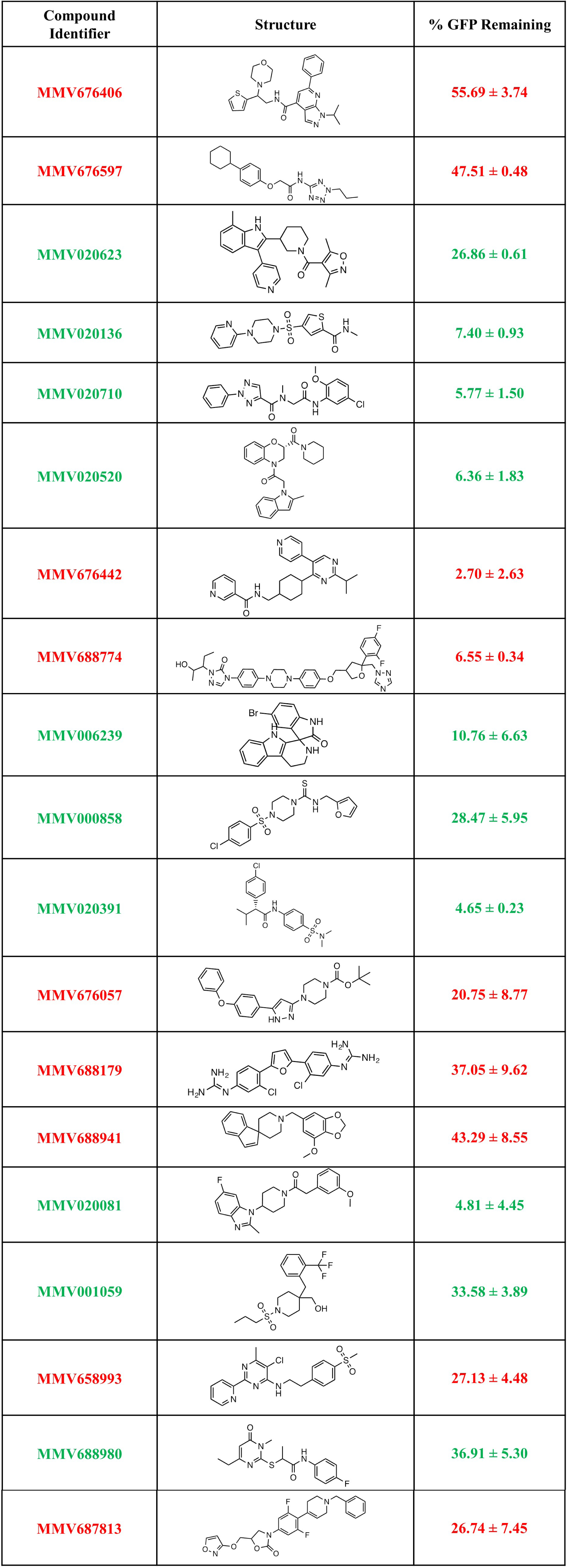
Candidates from the Pathogen Box disrupting lipid homeostasis. Compounds from the Pathogen Box that were identified as candidates inducing saponin sensitivity into the parasite plasma membrane (mean ± SEM from 2 independent experiments). Compounds in Green represent Na^+^-dependent and Red represent Na^+^- independent saponin sensitivity.

| Compound Identifier | Structure | % GFP Remaining |
| --- | --- | --- |
| **MMV676406** |  | **55.69 ± 3.74** |
| **MMV676597** |  | **47.51 ± 0.48** |
| **MMV020623** |  | **26.86 ± 0.61** |
| **MMV020136** |  | **7.40 ± 0.93** |
| **MMV020710** |  | **5.77 ± 1.50** |
| **MMV020520** |  | **6.36 ± 1.83** |
| **MMV676442** |  | **2.70 ± 2.63** |
| **MMV688774** |  | **6.55 ± 0.34** |
| **MMV006239** |  | **10.76 ± 6.63** |
| MMV000858 |  | 28.47 ± 5.95 |
| MMV020391 |  | 4.65 ± 0.23 |
| MMV676057 |  | 20.75 ± 8.77 |
| MMV688179 |  | 37.05 ± 9.62 |
| MMV688941 |  | 43.29 ± 8.55 |
| MMV020081 |  | 4.81 ± 4.45 |
| MMV001059 |  | 33.58 ± 3.89 |
| MMV658993 |  | 27.13 ± 4.48 |
| MMV688980 |  | 36.91 ± 5.30 |
| **MMV687813** |  | **26.74 ± 7.45** |

### Growth inhibition by Na^+^ influx independent compounds

The PfATP4 inhibitors such as pyrazoleamide and spiroindolones primarily cause parasite death by disrupting the Na^+^ homeostasis of the parasite ^7, 11^. The influx of Na^+^ into the parasite cytoplasm is followed by the altered lipid homeostasis of the parasite plasma membrane ^11^. We observed that the 50% inhibitory concentration of saponin sensitivity for the pyrazoleamides and spiroindolones was similar to their reported growth inhibitory EC_50_ (Figure 1B) ^4-5^. Since the Na^+^ influx-independent compounds targeted an unknown pathway, we evaluated those compounds for their ability to inhibit parasite growth and induce saponin sensitivity. We observed that the majority of compounds required micro-molar concentration in order to inhibit parasite growth, with the exception of MMV007113, MMV009108 and MMV688179 (Table 3; Column (+) PfNCR1). However, these compounds induced saponin sensitivity at 3-10 fold lower drug concentration, with the exception of MMV007907, MMV676057, MMV688179 and MMV687813 (Table 3). However, these 4 compounds did show heightened saponin sensitivity when the duration of treatment was increased to 4 h (Table 3). MMV007654, MMV019662 and MMV665908 were the most potent at disrupting plasma membrane lipid homeostasis and required 10-fold lower concentration for this effect compared to their growth inhibitory concentrations (Table 3), suggesting parasites can survive in vitro with abnormal distribution of PPM lipids. These results imply that the compounds identified in our screen primarily target the lipid homeostasis of the parasite.

**Table 3:** Activities of Na^+^ independent compounds. 11of 15 compounds identified were evaluated for their 50% saponin sensitivity inhibitory concentration (n=2). Growth inhibition was measured using ^3^H-hypoxanthine incorporation over 48 h treatment on parasites expressing PfNCR1 ((+) PfNCR1) or on parasites with PfNCR1 knocked-down ((-) PfNCR1) (n=3).

| Compound Identifier | Saponin Sensitivity IC<sub>50</sub>(μM) | Growth Inhibition EC<sub>50</sub> (μM) |  |
| --- | --- | --- | --- |
|  |  | (+) PfNCR1 | (-) PfNCR1 |
| MMV007113 | 0.66 ± .27 | 0.53 ± 0.09 | 0.49 ± 0.03 |
| MMV007907* | > 2 | 3.21 ± 0.30 | 0.16 ± 0.004 |
| MMV007654 | 0.16 ± 0.03 | 1.41 ± 0.07 | 0.04 ± 0.003 |
| MMV009108 | 0.14 ± 0.01 | 0.60 ± 0.03 | 0.02 ± 0.002 |
| MMV665908 | 0.10 ± 0.008 | 2.29 ± 0.08 | 0.06 ± 0.003 |
| MMV676442 | 0.56 ± 0.13 | 1.50 ± 0.12 | 0.04 ± 0.0003 |
| MMV688774 | 0.63 ± 0.03 | 3.74 ± 1.05 | 0.17 ± 0.03 |
| MMV676057* | 1.35 ± 0.57 | > 6 | 4.42 ± 0.4 |
| MMV688179* | > 2 | 0.66 ± 0.03 | 0.22 ± 0.006 |
| MMV687813* | > 2 | > 6 | 2.27 ± 0.1 |
| MMV019662 | 0.15 ± 0.03 | 1.68 ± 0.13 | - |

\* treatment for 4 h showed significantly lower GFP content after saponin exposure.

### Several distinct chemotypes target PfNCR1

Recently, a chemogenetic screen of the Malaria Box compounds found that resistance towards MMV009108 and MMV019662 required mutations in the gene PF3D7_0107500, now called Plasmodium falciparum Niemann-Pick Type C1-Related Protein (PfNCR1) ^12-13^. MMV009108 was able to reversibly induce saponin sensitivity without disrupting parasite Na^+^ homeostasis ^13^. Furthermore, we showed that MMV009108 and MMV019662 caused parasite death via the inhibition of PfNCR1 ^13^. Since these compounds were also identified in our current screen, we reasoned that the other candidates in our screen might also target PfNCR1. In order to test the specificity of the compounds towards PfNCR1 we employed a parasite line wherein expression of PfNCR1 was conditionally regulated. We utilized the TetR-DOZI-aptamer system in which expression of the targeted gene is dependent on anhydrotetracycline (aTc) ^19^. The withdrawal of aTc resulted in the efficient knockdown of PfNCR1 ^13^. Knock down parasites take several intraerythrocytic cycles to die, probably because some residual PfNCR1 protein allows them to survive for a limited time ^13^. We rationalized that parasites with PfNCR1 knocked-down should become hypersensitive towards compounds that target PfNCR1. We were able to show PfNCR1 knockdown parasites ((-) PfNCR1) became hypersensitive to the majority of compounds identified as Na^+^ independent inducers of saponin sensitivity ((+) PfNCR1 vs. (-) PfNCR1; Table 3). However, we also observed MMV007113 and MMV688179 did not appear to target PfNCR1. These compounds may target yet another component in the cholesterol maintenance pathway and therefore result in saponin sensitivity. In addition to MMV009108 and MMV019962, which were already shown to target PfNCR1, we have identified 5 new chemotypes that inhibit PfNCR1 resulting in the disruption of PPM lipid composition.

## Discussion

The Malaria and Pathogen Boxes were envisioned to further drug discovery and development by providing samples of chemically diverse compounds ^20^. In this study we have described a method to screen for compounds that target PPM lipid homeostasis. In accordance with our previous hypothesis ^11^ the Na^+^ influx (as determined by the Kirk group) correlated with saponin sensitivity for the majority of compounds. However, there were a few that did not correlate. We hypothesize that this discrepancy may be due to the differences in how the two screens were performed. The Na^+^ influx measurements were performed on saponin isolated parasites ^7, 15-16^, whereas saponin sensitivity assays were done with intact parasite infected RBCs. Saponin isolation leaves the host RBCs in a highly permeabilized state, devoid of hemoglobin and other soluble cytoplasmic proteins. This would allow the compounds direct access to the parasite plasma membrane. In contrast, the saponin sensitivity screen was performed on parasites that reside within an intact host cell. The presence of an intact RBC plasma membrane and cytoplasm could act as an impediment to some of the compounds resulting in the discrepancy we observed. This could also explain why Na^+^ influx occurred within seconds of treatment with compounds while saponin sensitivity was observed 30-45 min after treatment ^4, 7, 11^.

We identified 15 compounds that disrupted parasite lipid homeostasis without affecting the Na^+^ homeostasis of the parasite. We evaluated 11 of these compounds and found that their *in-vitro* growth inhibitory concentrations were >1μM. 7 of these compounds were extremely potent at inducing saponin sensitivity and appear to inhibit the function of PfNCR1. Since we observed that 4 h treatment with MMV007907, MMV676057, MMV688179 and MMV687813 resulted in a higher level of saponin sensitivity, there is a possibility that a longer duration of treatment may yield additional candidates from this screen.

Interestingly, we observed that MMV007113 and MMV676057 did not show any specificity towards PfNCR1, as assessed under PfNCR1 knockdown conditions. In mammals, cholesterol trafficking within the late endosome requires two indispensible proteins, NPC1 and NPC2 ^21-22^. NPC2 is thought to bind cholesterol released from lipoproteins within the endosome ^23^. NPC2 then transfers the cholesterol to the membrane bound NPC1, which transports the cholesterol out of the endosome ^23^. PfNCR1 could potentially function in a similar manner wherein one or several proteins participate in its function. PfNCR1 was localized to the plasma membrane of the parasite and could function as the sterol transporter in the membrane. Thus, MMV007113 and MMV676057 could potentially target a partner protein of PfNCR1 thereby resulting in saponin sensitivity independent of both PfNCR1 and PfATP4 inhibition.

The PfATP4 inhibitors like pyrazoleamide, spiroindolone and dihydroisoquinoline showed faster in vivo clearance compared to their *in vitro* killing rates ^4, 6, 24^. We have proposed that the incorporation of cholesterol into the parasite plasma membrane due to Na^+^ influx would lead to increased PPM rigidity ^11, 25^. The increased plasma membrane rigidity could result in restricted flow of infected RBCs through the host microvasculature and cause the parasites to be damaged under the extreme stress experienced in circulation ^26^. Cholesterol possibly enters the plasma membrane as the parasite ingests host hemoglobin. The uptake of hemoglobin by the parasite would lead to close proximity of the plasma membrane and the cholesterol-rich parasitophorous vacuolar membrane ^10^. A surveillance pathway would be needed ensure that the cholesterol does not traverse into the plasma membrane. Alternatively, this pathway may remove any cholesterol that diffuses into the plasma membrane. PfNCR1 has been implicated as the lipid transporter responsible for maintaining the plasma membrane lipid homeostasis ^13^. PfNCR1 may also serve to maintain sphingomyelin levels in the PPM similar to the function of *T. gondii* NCR1 in the inner membrane complex of *Toxoplasma* ^27^. Recently, both P*. falciparum* and *P. berghei* NCR1 deletions were shown to be lethal to the parasite ^13, 28^. This highlights the need for parasites to maintain appropriate cholesterol/lipid composition within the plasma membrane. PfNCR1inhibitors that were identified in our screen presented as relatively low potency anti-malarials *in vitro*. Nevertheless, these compounds were more potent at disrupting the lipid homeostasis of the parasite. We propose that compounds that disrupt parasite plasma membrane lipid homeostasis could serve as candidate antimalarials that clear parasites more efficiently *in vivo*. Our results indicate that PfNCR1 is inhibited by diverse chemotypes and may serve as the foundation for target based drug discovery.

## Material and Methods

### Cell Culture

*P. falciparum* NF54 yDHODH-GFP parasites were cultured in O^+^ human blood (Interstate Blood Bank, TN) in RPMI1640 supplemented with 15mM HEPES, 2g/L sodium bicarbonate, 10mg/L hypoxanthine, 50mg/L gentamicin sulfate, 0.5% Albumax II. Parasites were maintained at 5% hematocrit at 37°C in 90% N_2_, 5% CO_2_, 5% O_2_. All cultures were periodically tested to ensure they were free from mycoplasma contamination.

### Malaria Box and Pathogen Box

The Malaria Box and Pathogen Box were provided by Medicine for Malaria Venture (MMV) (Geneva, Switzerland). The compounds were provided at 10mM concentration in DMSO and were diluted into identical 96 well plates at 100μM in DMSO. The compound structures were available through ChEMBL (https://www.ebi.ac.uk/chembl/malaria/).

### Screening compounds and saponin treatment of parasites

Alanine synchronized trophozoite stage NF54 yDHODH-GFP parasites were plated in flat bottomed 96 well plates at 5% hematocrit. The compounds were added to a final concentration of 1μm ensuring 1%DMSO in all wells. DMSO was added to vehicle treated parasites at 1% v/v. The 96-well plates were placed at 37°C for 2 h in 90% N_2_, 5% CO_2_, 5% O_2_. The parasites were transferred from flat-bottomed culture plates to V-bottomed plates (NUNC) for saponin treatment and isolation. The parasite cultures were pelleted in V-bottomed plates (500g, 5 min) and parasite isolation was performed resuspending the parasite to 5% hematocrit in Albumax free RPMI. Saponin was added to the wells to a final concentration of 0.08% saponin w/v in Albumax-free RPMI and vigorously agitated by pipetting. An additional well with DMSO treated parasites was treated with 0.1% Triton X-100 to serve as an auto-fluorescence correction control. Isolated parasites were collected by centrifugation at 500g for 5 min and washed twice (500g, 5 min) with RPMI supplemented with Albumax to quench any remaining saponin. The parasites were lysed in radioimmunoprecipitation assay (RIPA) buffer (50mM Tris pH 7.4, 150mM NaCl, 1mM EDTA, 1% Triton X-100, 0.5% sodium deoxycholate) with protease inhibitor cocktail (Sigma-Aldrich). The lysates were transferred into flat-bottomed opaque 96- well plates (NUNC) and placed at 4°C for 30 min. Plates were centrifuged (500g, 15 min) to remove cell debris and plates were placed in a spectrofluorometer (Hitachi F-7000) to record GFP emissions (Ex:488nm; Em:510nm). The arbitrary fluorescent units were converted to % GFP remaining after auto-fluorescence correction using Microsoft Excel and plotted using Prism (Graphpad). Significant reduction in GFP content was assessed against 100nM Atv/1μM Pro treated parasites using a multiple t test (p < 0.05).

### Saponin sensitivity assays

To assess the saponin concentration used, synchronized trophozoite stage NF54 yDHODH-GFP parasites were incubated for 2 h with either DMSO or 100nM pyrazoleamide PA21A092 at 37°C in 90% N_2_, 5% CO_2_, 5% O_2_. Treated parasites were collected (500g, 5 min) and resuspended to 10% hematocrit in Albumax-free RPMI1640. Equal volumes of 2%, 0.66%, 0.11%, 0.074% and 0.024% saponin w/v were added to the resuspended parasites thereby making the final concentrations of saponin as 1%, 0.33%, 0.11%, 0.037% and 0.012% w/v. Saponin solutions were prepared as 3-fold serial dilution in Albumax-free RPMI1640 and maintained at 37°C. Parasites were agitated by repeated inversion for 10-15s, followed by centrifugation (500g, 5 min) Saponin treated parasites were washed twice with RPMI1640 w/ Albumax (500g, 5 min). Parasites were subsequently lysed in RIPA buffer (50mM Tris pH 7.4, 150mM NaCl, 1mM EDTA, 1% Triton X-100, 0.5% sodium deoxycholate) with protease inhibitors (Sigma-Aldrich). Lysed parasites were placed on ice for 30 min with intermittent agitation followed by centrifugation (14000g, 15 mins). The supernatant was transferred to 96well opaque plates (NUNC) and read in a Hitachi F-7000 fluorescence spectrophotometer (Ex: 488nm; Em: 510nm).

To determine saponin sensitivity IC_50_, synchronized trophozoite stage NF54 yDHODH-GFP parasites were incubated with the compounds at indicated concentrations for 2 h at 37°C in 90% N_2_, 5% CO_2_, 5% O_2_. The parasites were collected (500g, 5 min) and resuspended to 10% hematocrit in Albumax-free RPMI1640. Equal volumes of 0.16% w/v saponin prepared in Albumax-free RPMI (at 37°C) was added to the resuspended parasites resulting in a final saponin concentration of 0.08% saponin w/v. Parasites were agitated by repeated inversion for 10-15s, followed by centrifugation (500g, 5 min). Isolated parasites were washed in RPMI1640 supplemented with Albumax (500g, 5 min). Isolated parasites were lysed in RIPA buffer (50mM Tris pH 7.4, 150mM NaCl, 1mM EDTA, 1% Triton X-100, 0.5% sodium deoxycholate) with protease inhibitors (Calbiochem). Parasites were placed on ice for 30 min with intermittent agitation followed by centrifugation (14000g, 15 mins). The supernatant was transferred to 96well opaque plates (NUNC) and read in a Hitachi F-7000 fluorescence spectrophotometer (Ex: 488nm; Em: 510nm).

### PfNCR1 knockdown parasites

PfNCR1 knockdown parasites were generated as described by Istvan E, *et. al.*^13^. Briefly, a right homologous region (bp3671 - bp4893) and a left homologous region (a 948bp fragment starting 38bp past the stop codon) were amplified by PCR from gDNA and cloned into the vector pMG75. The gRNA Cas9 guide sequence (5’-TTAATGTAGTGGGCCAAAAC-3’) was cloned into the vector pyAIO. The guide and the homologous integration vectors were co-transfected and selected with 5μg/ml Blasticidin S. Parasites were cloned by limited dilution.

### Growth inhibition assay and PfNCR1 knockdown

PfNCR1 knockdown was performed by the withdrawing anhydrotetracycline (aTc). Synchronized trophozoite stage parasites were collected by centrifugation (500g, 5 min). Parasites were then resuspended to a final 20-25% hematocrit. Infected RBCs were separated from uninfected RBCs by layering onto top of 70%/35% Percoll cushion and centrifugation (500g, 20 min). Infected RBCs were collected at the 70%-35% Percoll interface and washed 3x with RPMI1640 (500g, 10 min). Parasites were then equally divided into culture flasks containing RBCs at 5% hematocrit in culture media with or without aTc. Parasites were placed at 37°C in 90% N_2_, 5% CO_2_, 5% O_2_ for 48 h and subsequently collected for assays.

Parasite growth inhibition was assessed using a modified version of previously described methods ^29^. Briefly, parasite growth inhibition was assessed by the incorporation of ^3^H-hypoxanthine. Synchronized parasites were diluted to 1% parasitemia at 3% hematocrit and treated at varying compound concentrations for 48 h. ^3^H-hyoxanthine incorporation was measured over the last 24 h. Plates were frozen at -80°C to stop growth and lyse cells. Lysate was transferred to EasyTabC glass fibre filters (Perkin Elmer) using Filtermate cell harvester and scintillation counts were measured using Microscint-O in Topcount NXT Beta Counter (Packard/ Perkin Elmer).

## Acknowledgements

We thank Dr. Hangjun Ke (Drexel University) for providing the NF54 yDHODH-GFP parasites. We also thank Dr. Eva S. Istvan (Washington University School of Medicine, St. Louis, MO) for kindly providing the parasites with regulatable PfNCR1. SB and ABV designed the experiments. SB, SN and JMM performed the experiments. SB, DEG and ABV wrote the manuscript with input from all authors.

## References

1. WHO, World Malaria Report. 2016.

2. Imwong, M.; Suwannasin, K.; Kunasol, C.; Sutawong, K.; Mayxay, M.; Rekol, H.; Smithuis, F. M.; Hlaing, T. M.; Tun, K. M.; van der Pluijm, R. W.; Tripura, R.; Miotto, O.; Menard, D.; Dhorda, M.; Day, N. P.; White, N. J.; Dondorp, A. M., The spread of artemisinin-resistant Plasmodium falciparum in the Greater Mekong Subregion: a molecular epidemiology observational study. Lancet Infect Dis 2017. DOI: 10.1016/S1473-3099(17)30048-8.

3. Lu, F.; Culleton, R.; Cao, J., Artemisinin-Resistant Plasmodium falciparum in Africa. N Engl J Med 2017, 377 (3), 306. DOI: 10.1056/NEJMc1705789.

4. Vaidya, A. B.; Morrisey, J. M.; Zhang, Z.; Das, S.; Daly, T. M.; Otto, T. D.; Spillman, N. J.; Wyvratt, M.; Siegl, P.; Marfurt, J.; Wirjanata, G.; Sebayang, B. F.; Price, R. N.; Chatterjee, A.; Nagle, A.; Stasiak, M.; Charman, S. A.; Angulo-Barturen, I.; Ferrer, S.; Belen Jimenez-Diaz, M.; Martinez, M. S.; Gamo, F. J.; Avery, V. M.; Ruecker, A.; Delves, M.; Kirk, K.; Berriman, M.; Kortagere, S.; Burrows, J.; Fan, E.; Bergman, L. W., Pyrazoleamide compounds are potent antimalarials that target Na+ homeostasis in intraerythrocytic Plasmodium falciparum. Nat Commun 2014, 5, 5521. DOI: 10.1038/ncomms6521.

5. Rottmann, M.; McNamara, C.; Yeung, B. K.; Lee, M. C.; Zou, B.; Russell, B.; Seitz, P.; Plouffe, D. M.; Dharia, N. V.; Tan, J.; Cohen, S. B.; Spencer, K. R.; Gonzalez-Paez, G. E.; Lakshminarayana, S. B.; Goh, A.; Suwanarusk, R.; Jegla, T.; Schmitt, E. K.; Beck, H. P.; Brun, R.; Nosten, F.; Renia, L.; Dartois, V.; Keller, T. H.; Fidock, D. A.; Winzeler, E. A.; Diagana, T. T., Spiroindolones, a potent compound class for the treatment of malaria. Science 2010, 329 (5996), 1175–80. DOI: 10.1126/science.1193225.

6. Jimenez-Diaz, M. B.; Ebert, D.; Salinas, Y.; Pradhan, A.; Lehane, A. M.; Myrand-Lapierre, M. E.; O’Loughlin, K. G.; Shackleford, D. M.; Justino de Almeida, M.; Carrillo, A. K.; Clark, J. A.; Dennis, A. S.; Diep, J.; Deng, X.; Duffy, S.; Endsley, A. N.; Fedewa, G.; Guiguemde, W. A.; Gomez, M. G.; Holbrook, G.; Horst, J.; Kim, C. C.; Liu, J.; Lee, M. C.; Matheny, A.; Martinez, M. S.; Miller, G.; Rodriguez-Alejandre, A.; Sanz, L.; Sigal, M.; Spillman, N. J.; Stein, P. D.; Wang, Z.; Zhu, F.; Waterson, D.; Knapp, S.; Shelat, A.; Avery, V. M.; Fidock, D. A.; Gamo, F. J.; Charman, S. A.; Mirsalis, J. C.; Ma, H.; Ferrer, S.; Kirk, K.; Angulo-Barturen, I.; Kyle, D. E.; DeRisi, J. L.; Floyd, D. M.; Guy, R. K., (+)-SJ733, a clinical candidate for malaria that acts through ATP4 to induce rapid host-mediated clearance of Plasmodium. Proc Natl Acad Sci U S A 2014, 111 (50), E5455–62. DOI: 10.1073/pnas.1414221111.

7. Spillman, N. J.; Allen, R. J.; McNamara, C. W.; Yeung, B. K.; Winzeler, E. A.; Diagana, T. T.; Kirk, K., Na(+) regulation in the malaria parasite Plasmodium falciparum involves the cation ATPase PfATP4 and is a target of the spiroindolone antimalarials. Cell Host Microbe 2013, 13 (2), 227–37. DOI: 10.1016/j.chom.2012.12.006.

8. Francis, G.; Kerem, Z.; Makkar, H. P.; Becker, K., The biological action of saponins in animal systems: a review. Br J Nutr 2002, 88 (6), 587–605. DOI: 10.1079/BJN2002725.

9. Lauer, S.; VanWye, J.; Harrison, T.; McManus, H.; Samuel, B. U.; Hiller, N. L.; Mohandas, N.; Haldar, K., Vacuolar uptake of host components, and a role for cholesterol and sphingomyelin in malarial infection. EMBO J 2000, 19 (14), 3556–64. DOI: 10.1093/emboj/19.14.3556.

10. Tokumasu, F.; Crivat, G.; Ackerman, H.; Hwang, J.; Wellems, T. E., Inward cholesterol gradient of the membrane system in P. falciparum-infected erythrocytes involves a dilution effect from parasite-produced lipids. Biol Open 2014, 3 (6), 529–41. DOI: 10.1242/bio.20147732.

11. Das, S.; Bhatanagar, S.; Morrisey, J. M.; Daly, T. M.; Burns, J. M., Jr.; Coppens, I.; Vaidya, A. B., Na+ Influx Induced by New Antimalarials Causes Rapid Alterations in the Cholesterol Content and Morphology of Plasmodium falciparum. PLoS Pathog 2016, 12 (5), e1005647. DOI: 10.1371/journal.ppat.1005647.

12. Cowell, A. N.; Istvan, E. S.; Lukens, A. K.; Gomez-Lorenzo, M. G.; Vanaerschot, M.; Sakata-Kato, T.; Flannery, E. L.; Magistrado, P.; Owen, E.; Abraham, M.; LaMonte, G.; Painter, H. J.; Williams, R. M.; Franco, V.; Linares, M.; Arriaga, I.; Bopp, S.; Corey, V. C.; Gnadig, N. F.; Coburn-Flynn, O.; Reimer, C.; Gupta, P.; Murithi, J. M.; Moura, P. A.; Fuchs, O.; Sasaki, E.; Kim, S. W.; Teng, C. H.; Wang, L. T.; Akidil, A.; Adjalley, S.; Willis, P. A.; Siegel, D.; Tanaseichuk, O.; Zhong, Y.; Zhou, Y.; Llinas, M.; Ottilie, S.; Gamo, F. J.; Lee, M. C. S.; Goldberg, D. E.; Fidock, D. A.; Wirth, D. F.; Winzeler, E. A., Mapping the malaria parasite druggable genome by using in vitro evolution and chemogenomics. Science 2018, 359 (6372), 191–199. DOI: 10.1126/science.aan4472.

13. Istvan, E. S.; Das, S.; Bhatnagar, S.; Beck, J. R.; Owen, E.; LLinas, M.; Ganesan, S. M.; Niles, J. C.; Winzeler, E. A.; Vaidya, A. B.; Goldberg, D. E., Plasmodium falciparum Niemann-Pick Type C1-Related Protein is a Druggable Target Required for Parasite Membrane Homeostasis. bioRxiv 2018. DOI: 10.1101/371484.

14. Spangenberg, T.; Burrows, J. N.; Kowalczyk, P.; McDonald, S.; Wells, T. N.; Willis, P., The open access malaria box: a drug discovery catalyst for neglected diseases. PLoS One 2013, 8 (6), e62906. DOI: 10.1371/journal.pone.0062906.

15. Lehane, A. M.; Ridgway, M. C.; Baker, E.; Kirk, K., Diverse chemotypes disrupt ion homeostasis in the Malaria parasite. Mol Microbiol 2014, 94 (2), 327–39. DOI: 10.1111/mmi.12765.

16. Dennis, A. S. M.; Rosling, J. E. O.; Lehane, A. M.; Kirk, K., Diverse antimalarials from whole-cell phenotypic screens disrupt malaria parasite ion and volume homeostasis. Sci Rep 2018, 8 (1), 8795. DOI: 10.1038/s41598-018-26819-1.

17. Painter, H. J.; Morrisey, J. M.; Mather, M. W.; Vaidya, A. B., Specific role of mitochondrial electron transport in blood-stage Plasmodium falciparum. Nature 2007, 446 (7131), 88–91. DOI: 10.1038/nature05572.

18. Ke, H.; Morrisey, J. M.; Ganesan, S. M.; Painter, H. J.; Mather, M. W.; Vaidya, A. B., Variation among Plasmodium falciparum strains in their reliance on mitochondrial electron transport chain function. Eukaryot Cell 2011, 10 (8), 1053–61. DOI: 10.1128/EC.05049-11.

19. Ganesan, S. M.; Falla, A.; Goldfless, S. J.; Nasamu, A. S.; Niles, J. C., Synthetic RNA-protein modules integrated with native translation mechanisms to control gene expression in malaria parasites. Nat Commun 2016, 7, 10727. DOI: 10.1038/ncomms10727.

20. Van Voorhis, W. C.; Adams, J. H.; Adelfio, R.; Ahyong, V.; Akabas, M. H.; Alano, P.; Alday, A.; Aleman Resto, Y.; Alsibaee, A.; Alzualde, A.; Andrews, K. T.; Avery, S. V.; Avery, V. M.; Ayong, L.; Baker, M.; Baker, S.; Ben Mamoun, C.; Bhatia, S.; Bickle, Q.; Bounaadja, L.; Bowling, T.; Bosch, J.; Boucher, L. E.; Boyom, F. F.; Brea, J.; Brennan, M.; Burton, A.; Caffrey, C. R.; Camarda, G.; Carrasquilla, M.; Carter, D.; Belen Cassera, M.; Chih-Chien Cheng, K.; Chindaudomsate, W.; Chubb, A.; Colon, B. L.; Colon-Lopez, D. D.; Corbett, Y.; Crowther, G. J.; Cowan, N.; D’Alessandro, S.; Le Dang, N.; Delves, M.; DeRisi, J. L.; Du, A. Y.; Duffy, S.; Abd El-Salam El-Sayed, S.; Ferdig, M. T.; Fernandez Robledo, J. A.; Fidock, D. A.; Florent, I.; Fokou, P. V.; Galstian, A.; Gamo, F. J.; Gokool, S.; Gold, B.; Golub, T.; Goldgof, G. M.; Guha, R.; Guiguemde, W. A.; Gural, N.; Guy, R. K.; Hansen, M. A.; Hanson, K. K.; Hemphill, A.; Hooft van Huijsduijnen, R.; Horii, T.; Horrocks, P.; Hughes, T. B.; Huston, C.; Igarashi, I.; Ingram-Sieber, K.; Itoe, M. A.; Jadhav, A.; Naranuntarat Jensen, A.; Jensen, L. T.; Jiang, R. H.; Kaiser, A.; Keiser, J.; Ketas, T.; Kicka, S.; Kim, S.; Kirk, K.; Kumar, V. P.; Kyle, D. E.; Lafuente, M. J.; Landfear, S.; Lee, N.; Lee, S.; Lehane, A. M.; Li, F.; Little, D.; Liu, L.; Llinas, M.; Loza, M. I.; Lubar, A.; Lucantoni, L.; Lucet, I.; Maes, L.; Mancama, D.; Mansour, N. R.; March, S.; McGowan, S.; Medina Vera, I.; Meister, S.; Mercer, L.; Mestres, J.; Mfopa, A. N.; Misra, R. N.; Moon, S.; Moore, J. P.; Morais Rodrigues da Costa, F.; Muller, J.; Muriana, A.; Nakazawa Hewitt, S.; Nare, B.; Nathan, C.; Narraidoo, N.; Nawaratna, S.; Ojo, K. K.; Ortiz, D.; Panic, G.; Papadatos, G.; Parapini, S.; Patra, K.; Pham, N.; Prats, S.; Plouffe, D. M.; Poulsen, S. A.; Pradhan, A.; Quevedo, C.; Quinn, R. J.; Rice, C. A.; Abdo Rizk, M.; Ruecker, A.; St Onge, R.; Salgado Ferreira, R.; Samra, J.; Robinett, N. G.; Schlecht, U.; Schmitt, M.; Silva Villela, F.; Silvestrini, F.; Sinden, R.; Smith, D. A.; Soldati, T.; Spitzmuller, A.; Stamm, S. M.; Sullivan, D. J.; Sullivan, W.; Suresh, S.; Suzuki, B. M.; Suzuki, Y.; Swamidass, S. J.; Taramelli, D.; Tchokouaha, L. R.; Theron, A.; Thomas, D.; Tonissen, K. F.; Townson, S.; Tripathi, A. K.; Trofimov, V.; Udenze, K. O.; Ullah, I.; Vallieres, C.; Vigil, E.; Vinetz, J. M.; Voong Vinh, P.; Vu, H.; Watanabe, N. A.; Weatherby, K.; White, P. M.; Wilks, A. F.; Winzeler, E. A.; Wojcik, E.; Wree, M.; Wu, W.; Yokoyama, N.; Zollo, P. H.; Abla, N.; Blasco, B.; Burrows, J.; Laleu, B.; Leroy, D.; Spangenberg, T.; Wells, T.; Willis, P. A., Open Source Drug Discovery with the Malaria Box Compound Collection for Neglected Diseases and Beyond. PLoS Pathog 2016, 12 (7), e1005763. DOI: 10.1371/journal.ppat.1005763.

21. Subramanian, K.; Balch, W. E., NPC1/NPC2 function as a tag team duo to mobilize cholesterol. Proc Natl Acad Sci U S A 2008, 105 (40), 15223–4. DOI: 10.1073/pnas.0808256105.

22. Luo, J.; Jiang, L.; Yang, H.; Song, B. L., Routes and mechanisms of post-endosomal cholesterol trafficking: A story that never ends. Traffic 2017, 18 (4), 209–217. DOI: 10.1111/tra.12471.

23. Ikonen, E., Cellular cholesterol trafficking and compartmentalization. Nat Rev Mol Cell Biol 2008, 9 (2), 125–38. DOI: 10.1038/nrm2336.

24. White, N. J.; Pukrittayakamee, S.; Phyo, A. P.; Rueangweerayut, R.; Nosten, F.; Jittamala, P.; Jeeyapant, A.; Jain, J. P.; Lefevre, G.; Li, R.; Magnusson, B.; Diagana, T. T.; Leong, F. J., Spiroindolone KAE609 for falciparum and vivax malaria. N Engl J Med 2014, 371 (5), 403–10. DOI: 10.1056/NEJMoa1315860.

25. Huang, J.; Feigenson, G. W., A microscopic interaction model of maximum solubility of cholesterol in lipid bilayers. Biophys J 1999, 76 (4), 2142–57. DOI: 10.1016/S0006-3495(99)77369-8.

26. Sui, B.; Gao, P.; Lin, Y.; Gao, B.; Liu, L.; An, J., Blood flow pattern and wall shear stress in the internal carotid arteries of healthy subjects. Acta Radiol 2008, 49 (7), 806–14. DOI: 10.1080/02841850802068624.

27. Lige, B.; Romano, J. D.; Bandaru, V. V.; Ehrenman, K.; Levitskaya, J.; Sampels, V.; Haughey, N. J.; Coppens, I., Deficiency of a Niemann-Pick, type C1-related protein in toxoplasma is associated with multiple lipidoses and increased pathogenicity. PLoS Pathog 2011, 7 (12), e1002410. DOI: 10.1371/journal.ppat.1002410.

28. Bushell, E.; Gomes, A. R.; Sanderson, T.; Anar, B.; Girling, G.; Herd, C.; Metcalf, T.; Modrzynska, K.; Schwach, F.; Martin, R. E.; Mather, M. W.; McFadden, G. I.; Parts, L.; Rutledge, G. G.; Vaidya, A. B.; Wengelnik, K.; Rayner, J. C.; Billker, O., Functional Profiling of a Plasmodium Genome Reveals an Abundance of Essential Genes. Cell 2017, 170 (2), 260–272 e8. DOI: 10.1016/j.cell.2017.06.030.

29. Desjardins, R. E.; Canfield, C. J.; Haynes, J. D.; Chulay, J. D., Quantitative assessment of antimalarial activity in vitro by a semiautomated microdilution technique. Antimicrob Agents Chemother 1979, 16 (6), 710–8.

